# Heritable endogenization of an RNA virus in a mammalian species

**DOI:** 10.1101/2020.01.19.911933

**Authors:** Atsuo Iida, Ronald Tarigan, Hiroshi Shimoda, Kanako Endo, Masaya Furuta, Hitoshi Takemae, Daisuke Hayasaka, Kenji Ichiyanagi, Ken Maeda, Eiichi Hondo

## Abstract

Viruses are considered one of the driving forces for genome rearrangements via infection and endogenization into the host genome (Kazazian, 2004; Maksakova et al., 2006). In 2010, Horie and colleagues demonstrated that a non-retroviral RNA virus, Borna disease virus (BDV), could integrate into the genome in cultured somatic cells (Horie et al., 2010). However, germline transmission of viral-derived sequences using animal models is yet to be experimentally demonstrated. In this study, we reported a case of heritable endogenization using the encephalomyocarditis virus (EMCV) and laboratory mice. The EMCV is a small non-enveloped single-strand RNA virus without its own reverse transcriptase activity (Carocci et al., 2012). Here, we demonstrated that the EMCV genomic RNA was reverse transcribed into DNA fragments in the murine testes. The DNA sequence originated from the RNA genome of EMCV was also detected in the liver and earlobes of the offspring generated from the EMCV-infected father. This suggests that the exogenous sequence derived from the EMCV is transmitted into the host germline and inherited across subsequent generations. This first experimental demonstration of viral endogenization proposes reconsideration about the impact of viruses as a driving force for genome modification.

## Main

Viruses are considered a significant threat to human society due to their high degree of virulence against the health of humans and domestic animals. Moreover, based on previous analyses of the retrovirus and ancestral records in the genome, viruses are considered a driving force for genomic rearrangement and diversification (Feschotte and Gilbert, 2012). It is a generally accepted hypothesis that viruses can integrate into the host genome, including the germline, becoming endogenous as a heritable factor (Stocking and Kozak, 2008). Some endogenous retroviral elements have the potential to place a variable mutational load by modifying their host genome (Kazazian, 2004; Maksakova et al., 2006). A previous study demonstrated that the Borna disease virus (BDV) integrates into the host genome using cultured mammalian cells (Horie et al., 2010). However, no study has experimentally demonstrated viral endogenization into a host’s germline and consequently becoming a heritable factor. Here, we demonstrate heritable viral transmission in a mammal using the encephalomyocarditis virus (EMCV), which is a non-enveloped single-strand RNA belonging to the Picornaviridae family (Figure 1A, Carocci et al., 2012). The EMCV is mainly a causative agent for heart failure involving myocarditis that can lead to death in several mammalian species (Hubbard et al., 1992; Papaioannou et al., 2003). Under laboratory conditions, various pathologies were reported in rodents by the artificial infection assay (Doi, 2011). In this study, we used the EMCV and laboratory mice (*Mus musculus*) as an experimental model to demonstrate viral endogenization into the host genome.

**Figure 1.**
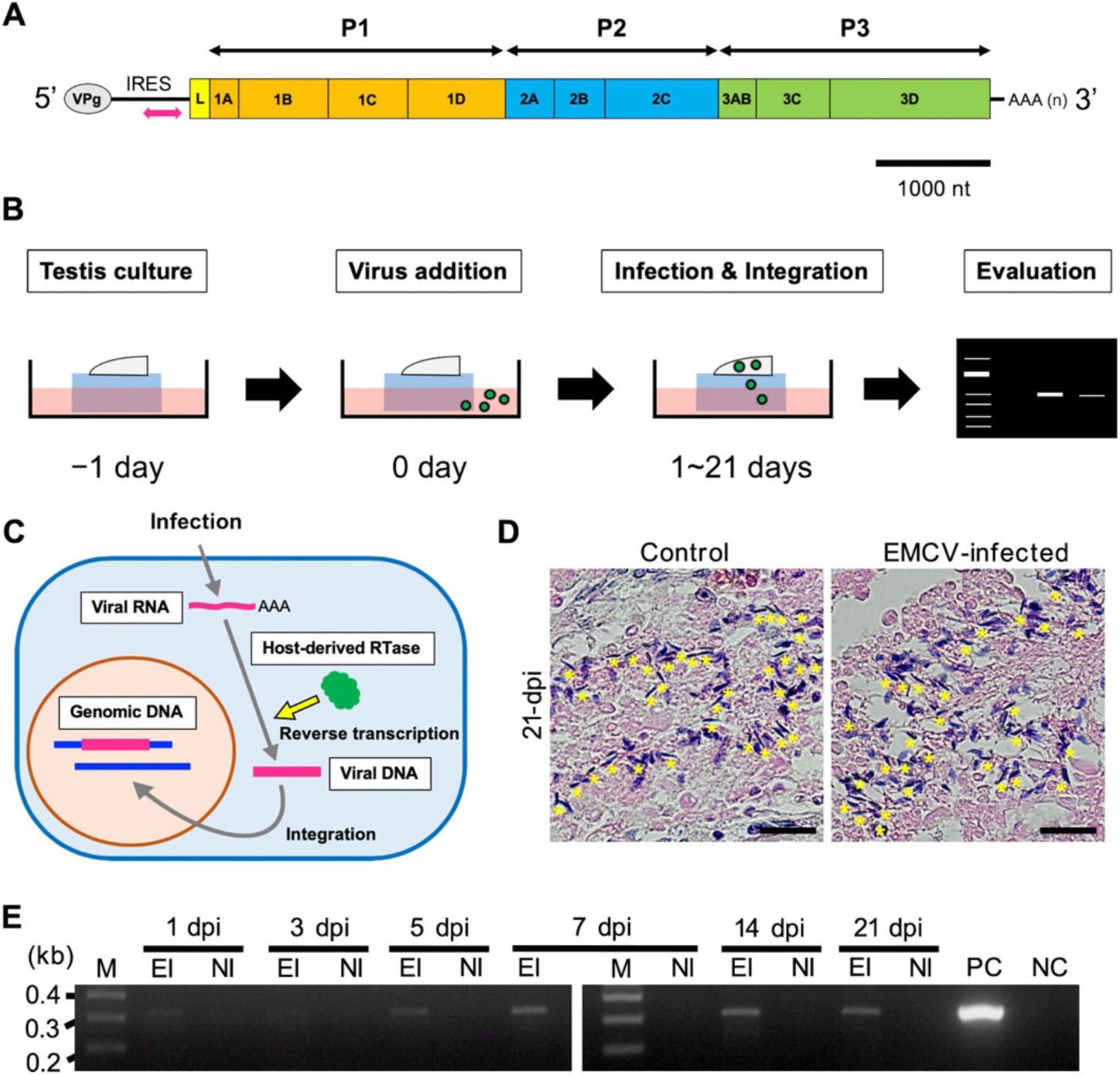
Reverse transcription of EMCV in cultured murine testis. **A**. Typical structure for the genomic RNA of EMCV. The magenta double head arrow indicates the PCR target region to detect a reverse transcribed EMCV genome. **B**. The illustration shows a strategy of the testis culture and viral infection. A reverse-transcription of the viral genomic RNA is evaluated by PCR and electrophoresis. **C**. A model for the pathway from viral infection to endogenization. A genomic sequence of an RNA virus is considered to be reverse-transcribed by host-derived reverse-transcriptase. Subsequently, the cDNA might be integrated into the host genome by non-homologous end joining. **D**. The photographs show typical images for transverse section of seminiferous tubule in the cultured testes at 21 dpi and the uninfected control. Arrowheads indicate spermatogonia. Scale bars, 20 µm. **E**. The electrophoresis shows a result of PCR to detect DNA fragment that is identical to the EMCV genome. The samples were harvested from the EMCV-infected testes at 1-, 3-, 5-, 7-, 14-, and 21-dpi. Expected size of the amplicon is 315 bp. M, size marker. EI, EMCV-infected. UI, uninfected. PC, positive control (EMCV cDNA). NC, negative control (empty vector).

### Viral infection into cultured testis

Mice testis can be cultured for over two months on agarose gel (Sato et al., 2011). To evaluate the viral infection and reverse transcription of the EMCV into the mammalian germline, we performed viral infection assay using the testis culture system. The EMCV strain NIID-NU1 was infected into the cultured testis extracted from the inbred C57BL/6JJmsSlc strain. Subsequently, spermatogenesis and reverse transcription of the viral genome were evaluated (Figure 1B). The EMCV is a non-retroviral RNA virus without its reverse-transcriptase (Carocci et al., 2012). Thus, if the viral-derived DNA was detected in the EMCV-infected testes, it confirms that the genomic RNA was reverse transcribed in the mammalian testes by the host-derived enzyme (Figure 1C). A histological observation indicated that the EMCV-infected testes were still alive until 30 days-post-infection (dpi), and spermatogenesis was maintained even up to 21 dpi (Figure 1D). The PCR amplicon corresponding to the 5’-untranslated region (5’-UTR) of the EMCV was detected in viral-infected testis cell lysates by PCR without reverse transcription (Figure 1E). This indicates that the EMCV genomic RNA can be reverse transcribed by host-derived factors. However, this observation of reverse transcription of the infected viral RNA has never been detected in cases of the Japanese encephalitis virus (JEV) (Figure S1). The host-derived reverse transcriptase might possess substrate specificity. As one of the trials investigating a responsible enzyme for reverse transcription of the EMCV genome into the murine genome, we focused on a telomerase reverse transcriptase (TERT) as a potential candidate involved in host-derived reverse transcription of the EMCV infectious viral particles (Nakamura et al., 1997; Weinrich et al., 1997). However, the EMCV DNA was also detected in the EMCV-infected cultured testes extracted from the TERT (−/−) mice (Figure S2). This suggests that other host factors play a role in reverse transcriptase activity of exogenous EMCV genomic RNA; however, we have yet to confirm this observation. Using an *in vitro* assay, these results support the hypothesis that transfection of the EMCV in gonadal cells integrates into the host germline, consequently becoming a viral genetic determinant in the offspring.

### Virus infection into an animal model

If the RNA virus plays a role in genomic rearrangements, the viral sequence should integrate into the host genome. To validate the endogenization *in vivo* and hereditability of viral-derived sequences, inbred C57BL/6JJmsSlc strain males were infected with the EMCV by intraperitoneal injection (Figure 2A, see also Figure S3). The EMCV exhibited high pathogenicity for the strain; thus, most of the injected mice became decayed and dead within 3 dpi (Figure 2B). On the other hand, low titer (1.0 × 10^−1^ pfu) EMCV was not shown the severe lethality, furthermore the survived males were fertile. We harvested DNA samples from the testes of dead males within 24 h after death. The amplicons signals for the 5’-UTR of the EMCV were obtained from the dead samples (Figure 2C). The amplicon was also obtained from a mating plug derived from the surviving male (Figure 2D). The plug sampling was performed at 46-50 dpi, when live EMCV was undetected from secretory fluids of the infected males (Figure S3). This strongly suggests that the EMCV genomic RNA is reverse-transcribed in the gonad including sperm, and subsequent integrations into the host genome might be occurred in these cells. Then, to verify a germline transmission and heredity of the integrated sequence, we obtained and analyzed the offspring mice from mating between the EMCV-infected males and uninfected females.

**Figure 2.**
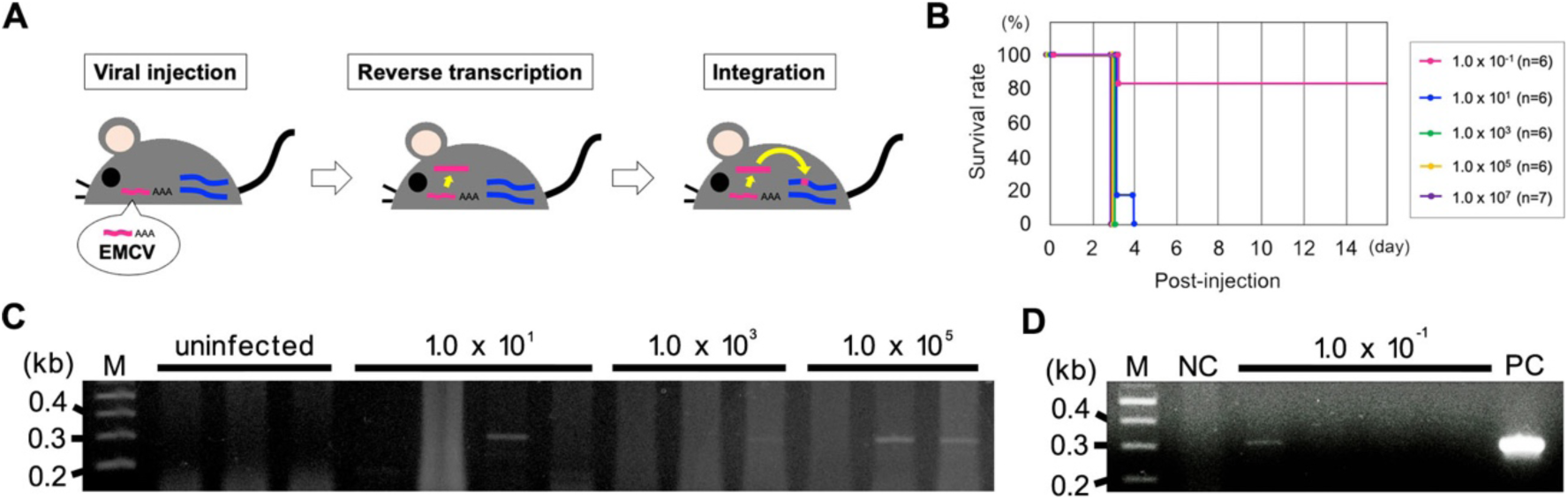
Reverse-transcription of EMCV in murine testis and mating plug. **A**. The illustration shows a strategy of the viral infection to live animal and a molecular hypothesis for subsequent reverse-transcription and endogenization. **B**. The line graph indicates a survival rate of the EMCV-infected adult male mice. All infected mice were dead within 4 dpi except lowest viral titer (1.0 × 10^−1^ pfu). **C**. The electrophoresis shows a result of PCR to detect DNA fragment that is identical to the EMCV genome. The DNA samples were harvested from testes extracted from the male mice deceased after EMCV injection at the titer of 10 × 10^1^, 10^3^, and 10^5^ pfu. Expected size of the amplicon is 315 bp. M, size marker. **D**. The electrophoresis shows a result of PCR to detect DNA fragment that is identical to the EMCV genome. The DNA samples were harvested from mating plugs derived from the male mice survived after EMCV injection at the titer of 10 × 10^−1^ pfu. Expected size of the amplicon is 315 bp. M, size marker. NC, negative control (empty vector). PC, positive control (EMCV cDNA).

### Viral integration and heredity

To validate the hereditability of the EMCV-derived sequence from the infected parent to the next generation, we obtained F1 offspring derived from the surviving males (Figure 3A, see also Figure S3). Approximately one-third of the offspring were dead within 3 d after birth, while the remaining offspring survived and grew normally (Figure 3B and S4). We harvested DNA from the liver of the dead offspring. The amplicon for the 5’-UTR of the EMCV was obtained from approximately half of the deceased offspring (Figure 3C). Next, we harvested DNA from the earlobe of the surviving offspring and their parental males. The amplicon was not detected from the parental male (Figure 3D). This suggests that no infected EMCV moved from the abdominal cavity to the earlobe and/or reverse transcribed in ear tissues.

**Figure 3.**
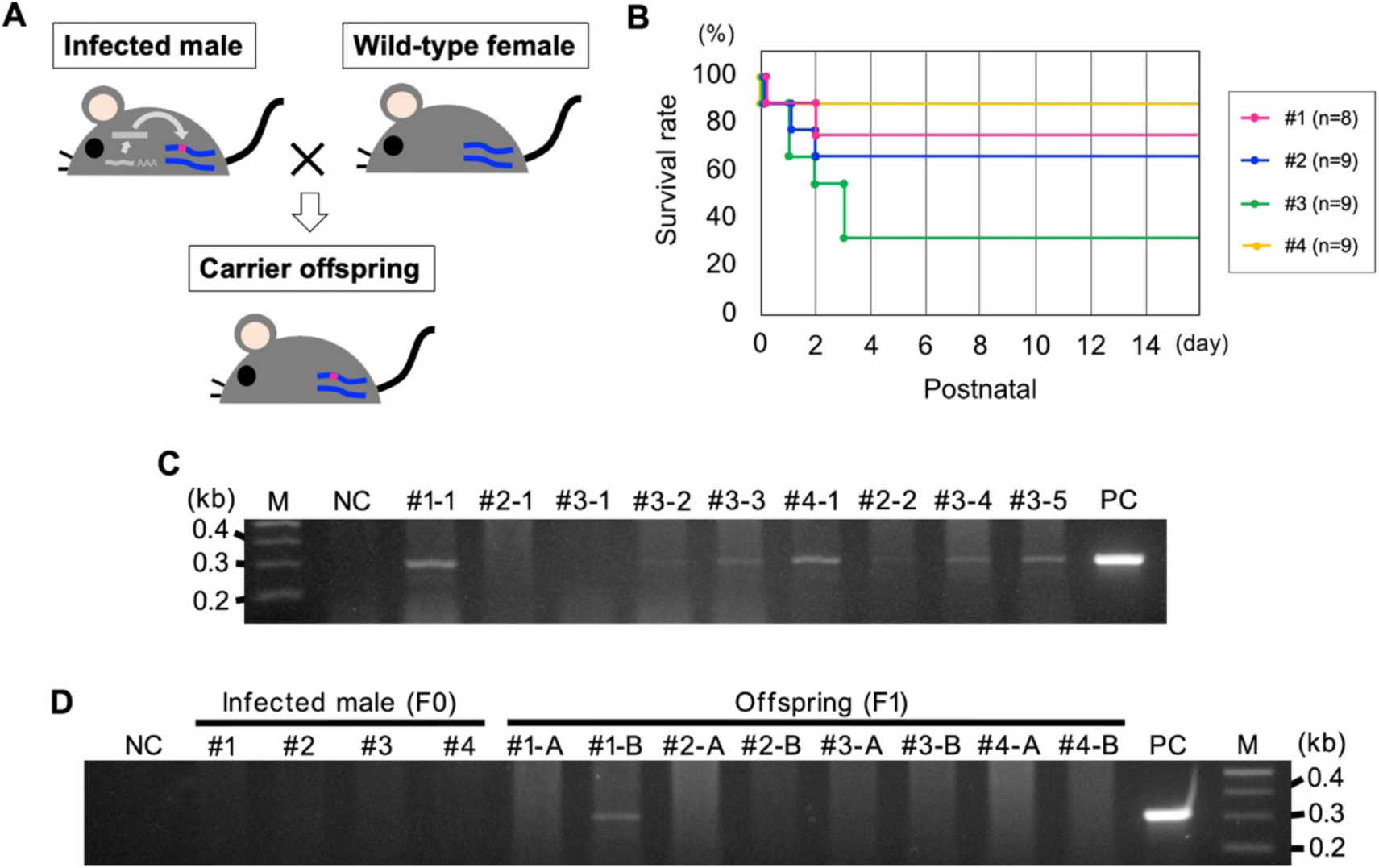
Heredity of EMCV-derived sequence from infected parent to offspring. **A**. The illustration shows a strategy to validate a heredity of the endogenized EMCV sequence from the infected male to the offspring. **B**. The line graph indicates a survival rate of F1 offspring obtained from the mating between the survived EMCV-infected male and uninfected female. Each color means the littermate derived from respective male mice. **C**. The electrophoresis shows a result of PCR to detect an endogenized EMCV fragment in the lethal offspring. The DNA samples were harvested from livers extracted from the F1 neonate deceased within 3 d after birth. The numbers were assigned for the parental male (#1∼#3) and littermate recognition. Expected size of the amplicon is 315 bp. M, size marker. NC, negative control (empty vector). PC, positive control (EMCV cDNA). **D**. The electrophoresis shows a result of PCR to detect an endogenized EMCV fragment in the survived offspring. The DNA samples were harvested from earlobes cut from the F1 offspring and their parent males. The numbers were assigned for the parental male (#1∼#4) and littermate recognition. Expected size of the amplicon is 315 bp. NC, negative control (empty vector). PC, positive control (EMCV cDNA). M, size marker.

Nonetheless, the amplicon was obtained from the earlobe of F1 offspring. This suggests that the reverse transcribed 5’-UTR of the EMCV integrates into the gametes of the infected male and is genetically inherited to the offspring. In this study, we checked only a small region corresponding to the 5’-UTR of the EMCV. Thus, it remains a possibility that we missed other novel integrations consisting of other regions of the EMCV. For a whole-genome survey, we are currently performing a deeper next generation sequence (NGS) analysis.

## Discussion

In this study, we report a case of germline transmission and hereditability of an RNA virus-derived sequence in a mammalian model. To date, this is the first study to experimentally demonstrate viral endogenization, except for indirect evidence based on the ancestral record in the genome. The EMCV genomic RNA was reverse transcribed in the cultured testes and mice germ cells, although the EMCV do not possess their own machinery for reverse transcriptase activity. The reverse transcription of non-retroviral RNA in mammalian cells has also been reported in the BDV (Horie et al., 2010). Our results indicate that the TERT is not involved in the reverse transcription step. Thus, other endogenous elements such as LINE-1 might be a source for reverse transcriptase activity in order to synthesize the EMCV cDNA (Brouha et al., 2003; Zhang et al., 2004). Understanding the precise mechanism involved in this step will require further molecular-based analysis using the EMCV-infected mice and cultured cell lines.

Our main achievement is the verification of the endogenization of the non-retroviral RNA virus into the mammalian germline and its hereditability into the next generation. Some offspring derived from the infected mice grew up healthy and fertile; thus, the newly integrated elements would continue to reside in the genome hereafter. As a pilot assay of next generation sequencing, we identified two reads consisting of a part of the EMCV genome and the mouse genome (Figure S5). These implications argue that non-retroviral RNA viruses could be one of the driving forces for mutations and rearrangements of the mammalian genome not only in the past but still in the present (Belyi et al., 2010; Katzourakis and Gifford, 2010). The endogenized element might contribute to genome rearrangements, including the acquisition of novel traits into the host (Feschotte and Gilbert, 2012). While it is not statistical data, the offspring of the EMCV-infected male mice exhibited smaller bodyweight than the average of the background strain (Figure S6). To demonstrate a causal relationship between EMCV transfection and potential novel traits, including changes in body weight, the surviving paternal mice and their offspring should be investigated over an extended period, or facilitate further mutations by additional viral infection.

Various mammalian species are considered natural carriers of the EMCV (Helwig and Schmidt, 1945; Gainer, 1967; Reddacliff et al., 1997; Billinis, 2009; Canelli et al., 2010). Our results indicate that these natural viral reservoirs not only include the risk of cross-transmission into other species but also endogenization and the addition of potential mutations in infected animals and their offspring. This study proposes to reconsider an impact of the EMCV and other viruses’ infection as a driving force for modifications and diversification of the host genome not only in the past, but also in the future.

## Methods

### Animal experiments

This study was approved by the Ethics Review Board for Animal Experiments of Nagoya University and Yamaguchi University. We sacrificed live animals in minimal numbers under anesthesia according to institutional guidelines.

### Mice strains and maintenance

The inbred C57BL/6JJmsSlc strain was used for all experiments in this study. The strain was purchased from Japan SLC, Inc. (Hamamatsu, Japan). EMCV infection into cultured testes was performed in the culture room at Nagoya University that adheres biosafety level 2 (BSL 2). EMCV infection into mice and breeding were performed in the Advanced Research Center for Laboratory Animal Science (ARCLAS) at Yamaguchi University that adheres to biosafety level 2 (BSL 2). Tissue samples, including active viruses, were maintained and manipulated in the laboratory of BSL 2. The virus was inactivated by paraformaldehyde before the export of tissues from the BSL 2.

### Virus isolation

The EMCV strain NIID-NU1 (DDBJ, #LC508268) was propagated in Vero cells (JCRB9013) purchased from the Japanese Collection of Research Bioresources (JCRB; Ibaraki, Japan). The EMCV-infected cells were incubated in Dulbecco’s Modified Eagle’s Medium (DMEM) with 10% fetal bovine serum (FBS) at 37 °C. The supernatant, including the released viral particles, was collected for *in vitro* or *in vivo* infection assay and stored at -80 °C until used.

### Tissue culture and viral infection

The testes culture was performed according to a previous report (Sato et al., 2011). Briefly, the testes were harvested from the inbred C57BL/6JJmsSlc strain males aged more than 6 weeks. The testes were cut into approximately 2 mm fragments and placed on the 0.8 % agarose in dishes filled by αMEM with 10% KnockOut™ Serum Replacement (Thermo Fisher Scientific K.K., Tokyo, Japan). The EMCV was added into the medium 1 d after the seeding. The tissue samples were harvested at 1, 3, 5, 7, 14, and 21 dpi, and the nuclear acid component was extracted from each sample. The samples were used as templates for PCR to detect the EMCV cDNA.

### PCR

PCR used to detect the viral cDNA was performed using the GoTaq^®^ Green Master Mix (Promega, Madison, WI, USA) under the following conditions: 120 s at 95 °C (initial denaturation), followed by 35 cycles of 30 s at 95 °C, 30 s at 55 °C, and 60 s at 72 °C; and 300 s at 72 °C (final elongations). Primer sequences are as follows: EMCV-forward (5’)-TGA ATG TCG TGA AGG AAG CAG T-(3’), EMCV-reverse (5’)-ACC TCG ACT AAA CAC ATG T-(3’), JEV-forward (5’)-CGC CCA TCA CCA GTG CGA G-(3’), JEV-reverse (5’)-TTG ATA ACA TCC ACT TG-(3’).

### Viral infection into mice

The EMCV was injected into the abdominal cavity of 6-week-old inbred C57BL/6JJmsSlc strain males. The viral titers were 10^7^, 10^5^, 10^3^, 10, and 10^−1^ plaque-forming units (PFUs). The testes were extracted from the deceased mice within 24 hours after death. At day 26 post-infection, the surviving males were crossed with uninfected 12-week-old inbred C57BL/6JJmsSlc strain females. One male mouse was transferred into the cage with one female at 7 PM. Their plug was checked and harvested the following morning. After the birth of F1 mice, neonatal mice monitored daily for their health condition. At 69th day post-infection, earlobes from F1 mice and survived EMCV infected male mice were collected.

## Author contributions

E.H. designed the experiments. A.I., H.S., R.T., K.E., M.F., and H.T performed the experiments and analyses. H.D, K.I., and K.M. helped with the experimental design and analyses. A.I. and E.H. wrote the manuscript, and all authors approved the final version submitted.

## Supplementary figures

**Figure S1.**
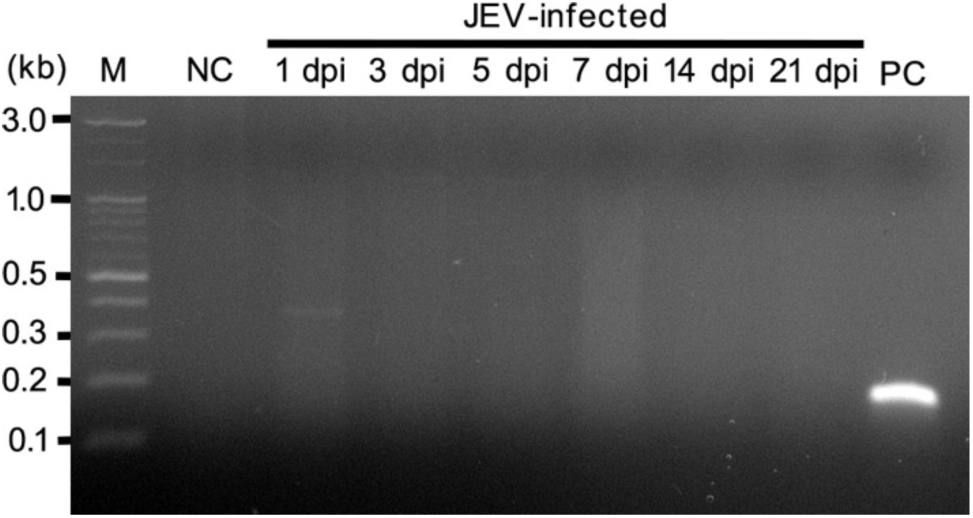
Detect for reverse-transcription of JEV in cultured murine testis. The electrophoresis shows a result of PCR to detect the DNA fragment that is identical to the JEV genome. The samples were harvested from the JEV-infected testes at 1-, 3-, 5-7-, 14-, and 21-dpi. Expected size of the amplicon is 161 bp. M, size marker. NC, negative control (empty vector). PC, positive control (JEV cDNA).

**Figure S2.**
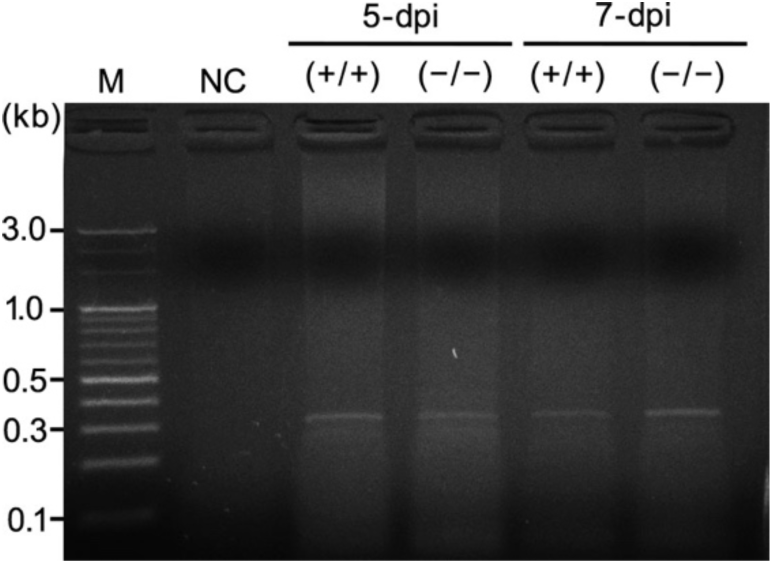
Detect for reverse-transcription of EMCV in TERT (−/−) cultured murine testis. The electrophoresis shows a result of PCR to detect DNA fragment that is identical to the EMCV genome. The samples were harvested from the EMCV-infected TERT (+/+) or TERT (−/−) littermate testes at 5- and 7-dpi. Expected size of the amplicon is 315 bp. M, size marker. NC, negative control (uninfected testis).

**Figure S3.**
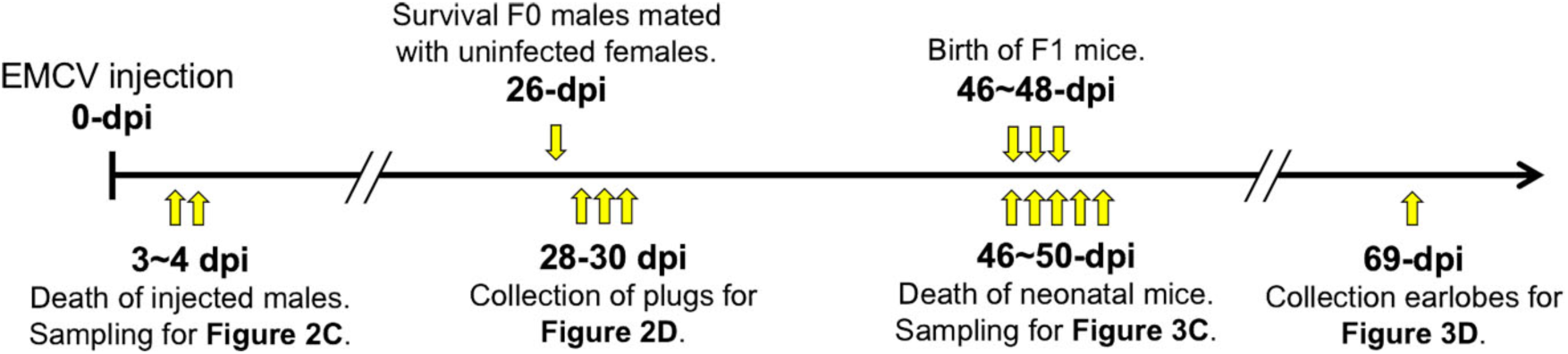
Experimental flow for viral infection to live animals. The illustration shows the time course of the EMCV infection to mice and the actual timing for the important events and sampling in this study. The events described above are about the infection and mating. The others are samplings mentioned in the main text and figures.

**Figure S4.**
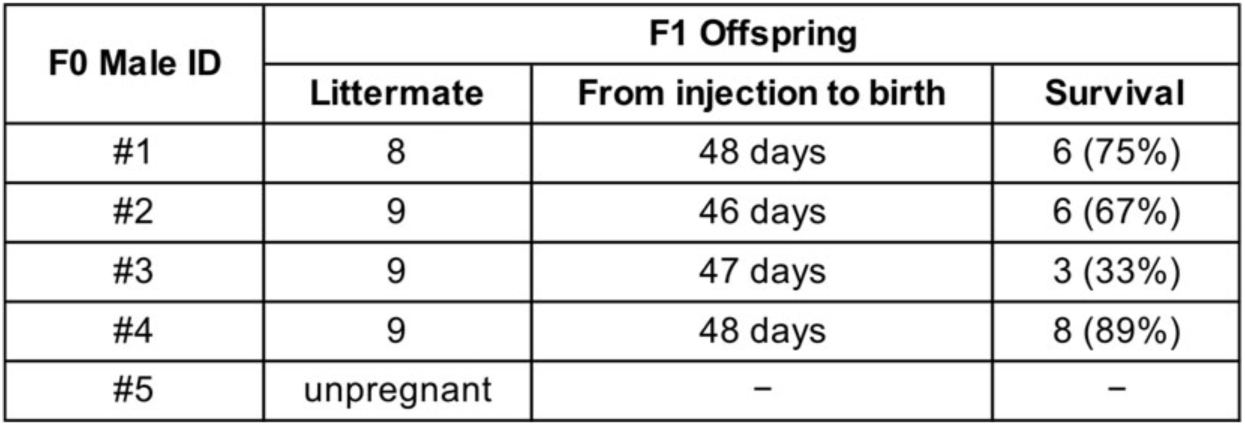
Numerical data for survival rate of the F1 generation mice. The table shows the numerical data for survival rate of the F1 generation mice obtained from the mating between the EMCV-infected male and uninfected female. This figure is related to Figure 3B. Survival number was recorded every day until 15 days after the birth.

**Figure S5.**
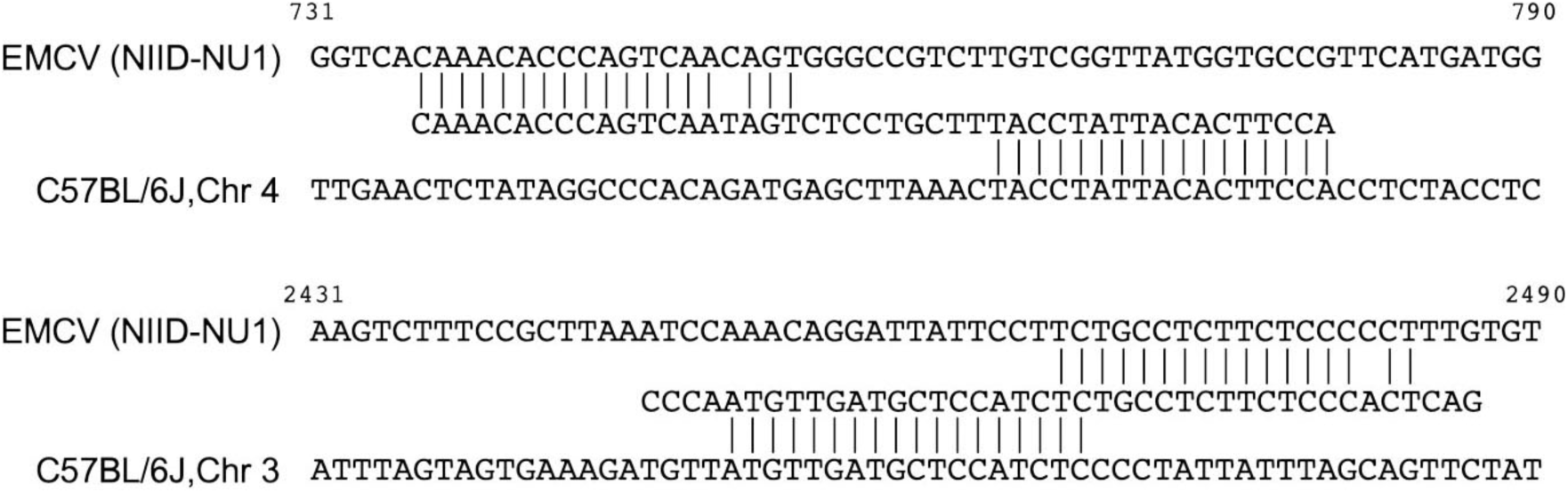
Sequences of the EMCV integration locus. The alignments illustration shows the raw sequence reads mapped to the EMCV and mouse genome. The genomic DNA extracted from the EMCV-infected testes was used for the NGS. The above read contains a small insertion in the border of the EMCV and mouse sequences. The other is considered to have undergone a precise non-homologous end joining, while an extra sequence was observed in the terminals of the read.

**Figure S6.**
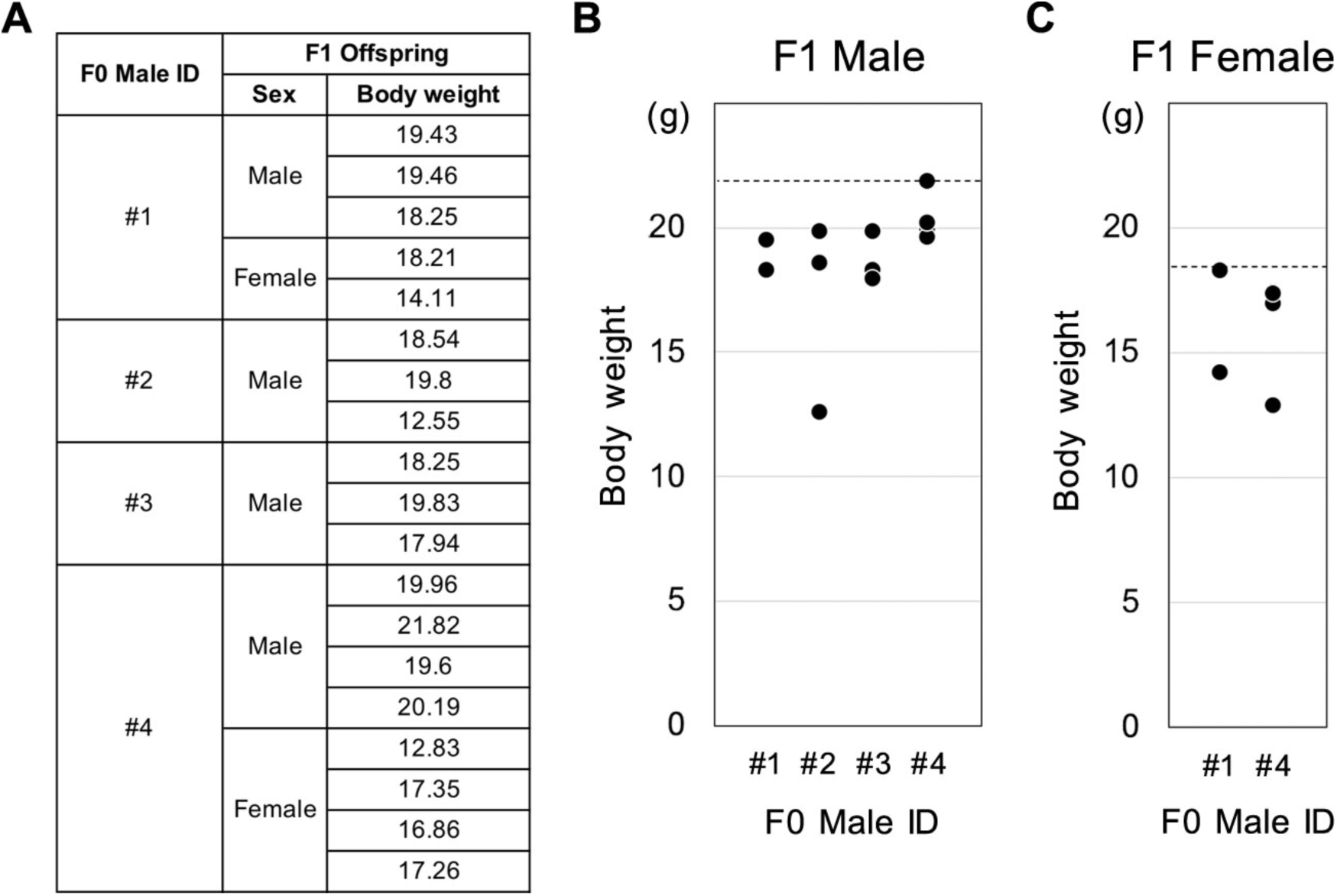
Body weight of the survived F1 generation mice. **A-C.** The table and graphs show the body weight of the survived F1 generation mice measured at 6-weeks. The offspring of the EMCV-infected male mice tended to exhibit smaller weight than the average of the C57BL/6J strain (Male, 21.9 ± 1.8. Female, 18.5 ± 0.9.). The dotted lines in **B** and **C** indicate the averages in each sex. The data were obtained from the Jackson Laboratory website (https://www.jax.org/jax-mice-and-services/strain-data-sheet-pages/body-weight-chart-000664).

